# The Mammalian Target of Rapamycin (mTOR) Kinase Mediates Haloperidol-Induced Cataleptic Behavior

**DOI:** 10.1101/2020.06.18.160416

**Authors:** Uri Nimrod Ramírez-Jarquín, Neelam Shahani, William Pryor, Alessandro Usiello, Srinivasa Subramaniam

**Affiliations:** Department of Neuroscience, The Scripps Research Institute, Florida, Jupiter, Florida 33458; Department of Environmental, Biological, and Pharmaceutical Sciences and Technologies, University of Campania Luigi Vanvitelli, 81100, Caserta, Italy; Laboratory of Behavioral Neuroscience, Ceinge Biotecnologie Avanzate, 80145, Naples, Italy

**Keywords:** Psychosis, Freezing, Extra pyramidal symptoms, Pre-clinical model, Medium spiny neurons, Tissue-specific regulation, Brain behavior

## Abstract

The mammalian target of rapamycin (mTOR) is a ubiquitously expressed serine/threonine kinase protein complex (mTORC1 or mTORC2) that orchestrates diverse functions ranging from embryonic development to aging. However, its brain tissue-specific roles remain less explored. Here, we have identified that the depletion of the *mTOR* gene in the mice striatum completely prevented the extrapyramidal motor side-effects (catalepsy) induced by the dopamine 2 receptor (D2R) antagonist haloperidol, which is the most widely used typical antipsychotic drug. Conversely, a lack of striatal mTOR in mice did not affect catalepsy triggered by the dopamine 1 receptor (D1R) antagonist SCH23390. Along with the lack of cataleptic effects, the administration of haloperidol in mTOR mutants failed to increase striatal phosphorylation levels of ribosomal protein pS6 (S235/236) as seen in control animals. To confirm the observations of the genetic approach, we used a pharmacological method and determined that the mTORC1 inhibitor rapamycin has a profound influence upon post-synaptic D2R-dependent functions. We consistently found that pretreatment with rapamycin entirely prevented (in a time-dependent manner) the haloperidol-induced catalepsy in wild-type mice. Collectively, our data indicate that striatal mTORC1 blockade may offer therapeutic benefits with regard to the prevention of D2R-dependent extrapyramidal motor side-effects of haloperidol in psychiatric illness.

## INTRODUCTION

mTOR exists as the mTOR-mLST8-Raptor complex (mTORC1) and mTOR-mLST8-Rictor complex (mTORC2). It serves as a multifunctional kinase in embryonic development, cancer, diabetes, aging, and neurodegenerative diseases (Bockaert and Marin, 2015; Laplante and Sabatini, 2012; Stallone et al., 2019). Its role and regulation in nervous system physiology and disease, however, is poorly understood (Hoeffer and Klann, 2010). This represents a major knowledge gap because the malfunction of mTORC1 activity (either by being too high or too low) has been linked to a variety of brain dysfunctions such as epilepsy, mental retardation, tuberous sclerosis, Huntington disease (HD), Parkinson’s disease (PD), and Alzheimer’s disease (AD), all of which affect a specific set of neuronal populations in the brain (Caccamo et al., 2014; Malagelada et al., 2010; Ravikumar et al., 2004; Troca-Marin et al., 2012; Zeng et al., 2009). A detailed understanding of how mTOR is regulated and what role it plays in selective brain regions is important for the development of better intervention strategies.

The brain’s striatum is composed of more than 95% inhibitory medium spiny neurons (MSNs) and it plays an important role in motor, cognitive, psychiatric, and reward behaviors (Grahn et al., 2008). MSNs dysfunctions can lead to the motor abnormalities seen in HD and PD; however, the molecular mechanisms are unclear. Interestingly, global blocking of mTORC1 signaling with rapamycin affords protection against the pathological and behavioral symptoms associated with HD and PD in murine models (Crews et al., 2010; Dehay et al., 2010; Fox et al., 2010; Malagelada et al., 2010; Ravikumar et al., 2004; Sarkar et al., 2008). However, the striatal-specific roles of mTOR signaling remains obscure.

Two major types of functionally distinct MSN are recognized, based on the dopamine 1 receptor (D1R) or dopamine 2 receptor (D2R) expression found in the striatum. In association with other receptors (e.g., glutamate, serotonin, and adenosine A1 and A2A receptors), dopamine receptors play critical roles in the processing of sensory, motor, cognitive, and motivational functions. (Graybiel and Grafton, 2015; Rolls, 1994). Functionally, D1R signaling increases Gαolf/adenylyl cyclase/cAMP/PKA signaling in the direct pathway of the basal ganglia, whereas D2R signaling inhibits cAMP/PKA signaling in the indirect pathway (Fernandez-Duenas et al., 2019; Herve, 2011; Kuroiwa et al., 2012; Nishi et al., 2011). Both dopamine D1 and D2 receptor stimulation promote motor activity. Pharmacological inhibition either of D1R or the D2R consistently trigger severe motor deficit and extrapyramidal side effects (EPS) (Klemm, 1989).

Recent studies have indicated that coordinated signaling of both D1R and D2R is responsible for the initiation and execution of motor activity (Sheng et al., 2019). Importantly, mTOR phosphorylation is selectively increased in the striatum during L-DOPA-induced dyskinesia (Eshraghi et al., 2020) and motor learning (Bergeron et al., 2014). However, the genetic evidence for the physiological role of mTOR signaling in the striatum (or its role in D1R versus D2R MSNs signaling) is currently unknown. Using genetic and pharmacological approaches, we investigated the role of mTOR on striatal-mediated motor behaviors under basal and challenged conditions.

## RESULTS

### Striatal mTOR regulates motor behaviors

The role of mTOR signaling in the regulation of striatal motor functions under basal conditions remains unclear. To address this question, we carried out conditional depletion of *mTOR* in the striatum of adult *mTOR^flox/flox^* mice. We used an AAV1.hSyn.HI.WPRE.SV40 variant expressing Cre-GFP (AAV-Cre-GFP) under the control of human synapsin promoter to deplete *mTOR* preferentially in striatal neuronal cells (Kugler et al., 2003). We stereotaxically injected purified virus (AAV-Cre-GFP or AAV-GFP) bilaterally into the striatum of 8-week-old *mTOR^flox/flox^* mice (Fig. 1A, B). Using Ctip2 (a marker for MSNs), we confirmed that in AAV-Cre-GFP-injected *mTOR^flox/flox^* mice (*mTOR* mutant), mTOR is depleted in the striatum 18 weeks after Cre injection, as expected, but not in AAV-GFP-injected *mTOR^flox/flox^* mice (control) (Fig. 1C, D, E). To determine the potential influence of neuronal *mTOR* depletion on cell survival, we estimated the number of cells and ventricular size between *mTOR* mutant and control mice. We found no gross changes in the number of total cells (Fig. 1E) or the ventricular size of the rostral and caudal striatal regions between *mTOR* mutant and control mice (Fig. 1F, G). These results indicate that AAV-Cre-GFP injection produces *mTOR* depletion in the MSNs and does not elicit any neurodegenerative-like phenotype.

**Figure 1.**
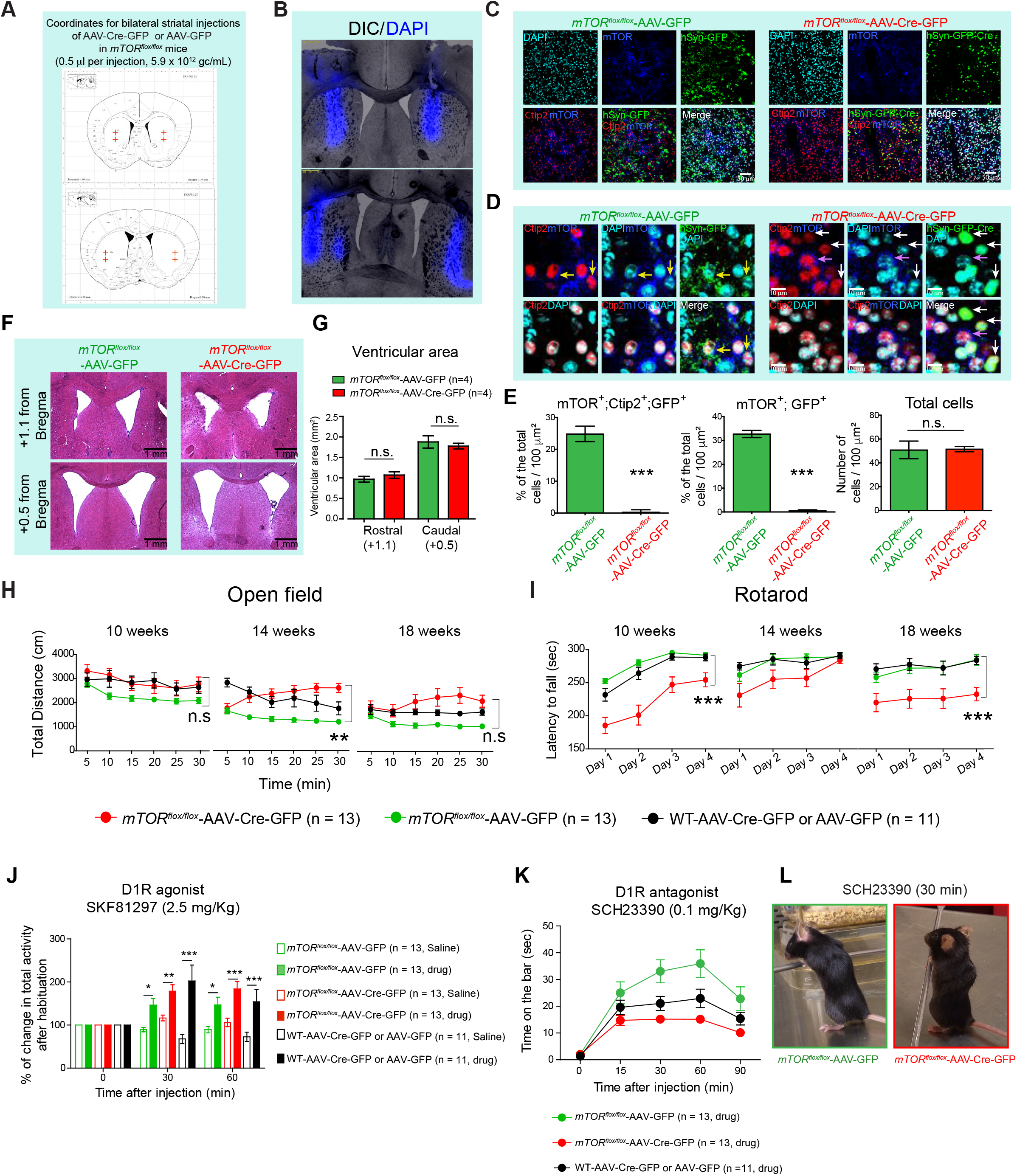
Effect of striatal mTOR depletion on motor behaviors. (**A**) Schematic representation of the AAV-Cre-GFP or AAV-GFP injected sites at the indicated coordinates targeting dorsal side of mice striatum. (**B**) Representative section showing the DAPI (blue) injection in the striatum using the coordinates in A. (**C**) Confocal images of the striatal brain sections from the *mTOR^flox/flox^* mice injected with AAV-Cre-GFP or AAV-GFP, showing GFP or GFP-Cre (green) expression, mTOR (blue), and Ctip2 (red) immunohistochemistry, and nuclear stain, DAPI (cyan). (**D**) High magnification of confocal images in C, showing that in AAV-GFP injected *mTOR^flox/flox^* mice, Ctip2-positive medium spiny neurons (MSNs) show GFP expression and mTOR immunostaining (yellow arrows). In AAV-Cre-GFP injected *mTOR^flox/flox^* mice, Ctip2 positive MSNs express GFP (cre) but are negative for mTOR immunostaining (white arrows). Some Ctip2 positive MSN negative for GFP (cre) are positive for mTOR staining (pink arrow). **(E**) Quantification for total number of cells identified by DAPI staining, % of mTOR, Ctip2 and GFP triple-positive neurons and % of mTOR and GFP double-positive neurons in striatum of the *mTOR^flox/flox^* mice injected with AAV-Cre-GFP or AAV-GFP. Images are representative of five ROIs from 4-5 sections per animal (n= 4 mice per group). Percentages were determined by considering the number of DAPI stained nuclei as 100%. All values are mean ± SEM. n.s. not significant, ***P < 0.001, two-tailed Student’s *t*-test. (**F**) Representative hematoxylin/eosin-stained sections for rostral (+1.1 from bregma) and caudal (+0.5 from bregma) ventriculus in *mTOR^flox/flox^* mice injected with AAV-Cre-GFP or AAV-GFP. (**G**) Quantification of ventricular area from F. n.s. not significant, two-way ANOVA, Bonferroni post-hoc test (four caudal and four rostral sections were quantified for four mice per group). (**H, I**) Total distance (cm) at the indicated time points in open-field test (OFT) (H) and latency to fall (sec) in rotarod test (I) for the *mTOR^flox/flox^* injected with AAV-GFP (n=13, female =10, male =3), AAV-Cre-GFP (n=13, female =6, male =7) or WT mice injected with AAV-GFP or AAV-Cre-GFP (n=11, female =5, male =6) at 10, 14 and 18 weeks of age. Data are mean ± SEM. **P < 0.01, ***P< 0.001, repeated measures two-way ANOVA followed by Bonferroni post-hoc test. (**J**) D1R agonist (SKF81297, 2.5 mg/Kg, i.p.)-induced activity in OFT in AAV-Cre-GFP or AAV-GFP injected *mTOR^foxlflox^* and AAV-Cre-GFP/GFP injected WT mice. Bar graphs indicates % of change in total activity after habituation. Data are mean ± SEM, n = 11-13 per group, *P <0.05, **P < 0.01, ***P < 0.001, repeated measures two-way ANOVA followed by Bonferroni post-hoc test. (**K**) Quantification of the catalepsy (time on the bar, sec)-induced by D1R antagonist SCH23390 (0.1 mg/Kg, i.p.) in indicated mice groups. Data are mean ± SEM, n = 11-13 per group, repeated measures two-way ANOVA followed by Bonferroni post-hoc test. (**L**) Representative image of catalepsy in AAV-Cre-GFP or AAV-GFP injected *mTOR^foxlflox^* mice treated with SCH23390.

We next assessed the striatal motor functions in *mTOR* mutant and control mice two weeks after AAV-Cre-GFP or AAV-GFP control injections. As Cre-recombinase injection in the brain may affect behaviors (Rezai Amin et al., 2019), we have included an additional control group for Cre: WT [C57BL/6] mice injected with AAV-Cre-GFP (Cre-control) or AAV-GFP (GFP-control) in all of our longitudinal behavioral analyses.

First, we measured locomotor activity using the open-field test (OFT). In the OFT, *mTOR* mutant, control, *and* Cre/GFP-control mice are placed individually in faintly lit open field chambers for 30 min sessions. The *mTOR* mutant mice displayed a mild increase in forward locomotion at 14 weeks that was not significantly different at 10 or 18 weeks of age, compared to Cre/GFP injected control animals (Fig. 1H).

Second, we investigated whether depletion of striatal mTOR impacts on balance and motor coordination, which is regulated by the striatum-cerebellar circuitry, using rotarod(Bostan and Strick, 2018). The *mTOR* mutant showed a decreased trend of motor coordination on the rotarod test compared to the control and Cre/GFP-control groups (Fig. 1I).

Overall, these results indicate that striatal mTOR plays a modulatory role in locomotion and motor coordination under basal conditions.

### Striatal mTOR does not influence D1R-mediated motor effects

Dopamine regulates motor functions such as locomotion and motor coordination by stimulating two main classes of receptors in the striatum (D1R and D2R) (Durieux et al., 2012). Considering that striatal mTOR depletion produces motor alterations under basal conditions, we questioned to what extent the D1R signaling-mediated function is affected in *mTOR* mutant mice. To address this question, we intraperitoneally (i.p.) injected pharmacological modulators that either activate D1R-signaling using SKF81297 (2.5 mg/kg, i.p.) or inhibit D1R-signaling using SCH23390 (0.1 mg/kg, i.p.), as described in previous studies (Ghiglieri et al., 2015; Napolitano et al., 2010; Usiello et al., 2000; Vitucci et al., 2016). Injection of SKF81297 (2.5 mg/kg, i.p.), a selective agonist of the D1R receptor, produced robust motor stimulation in all animals compared to the saline-administered group. Thus, administration of the D1R agonist induced comparable hyperlocomotion in AAV-Cre-GFP-injected *mTOR^flox/flox^* and control groups (*mTOR^flox/flox^* injected with AAV-GFP or WT mice injected with AAV-Cre-GFP or AAV-GFP) when tested at 30 and 60 min (Fig. 1J). This result indicates that mTOR depletion does not grossly interfere with D1R-mediated motor stimulation.

We next asked whether striatal mTOR plays any role in D1R-mediated catalepsy. Indeed, it has been well-established that blocking D1R with its antagonist SCH23390 (0.1 mg/kg, i.p.) elicits cataleptic behavior (i.e., the animal was unable to correct an externally imposed posture—time spent on the bar) (Morelli and Di Chiara, 1985; Napolitano et al., 2019). Notably, SCH23390 administration induced a similar cataleptic response in *mTOR* mutant mice and control groups (Fig. 1K, L). This result indicates that mTOR depletion has no significant effect on D1R antagonist-induced EPS.

Collectively, this data indicates that striatal mTOR does not affect pharmacologically modulated D1R-dependent motor behaviors.

### Striatal mTOR promotes D2R inhibition (Haloperidol)-induced cataleptic behavior

We then investigated whether mTOR depletion modulates pre-and post-synaptic D2R-signaling-mediated motor behavior. We administered quinpirole (0.5 mg/kg, i.p.), a D2R agonist, that by activating the presynaptic D2R reduces dopamine concentration in the striatum and in turn exerts overall dopamine receptor hypo-stimulation coupled to motor depression in mice(Napolitano et al., 2010; Radl et al., 2018; Usiello et al., 2000). Interestingly, we found that regardless of genotype, the administration of quinpirole similarly inhibited motor exploration in a novel environment (Fig. 2A). This data indicates that a lack of mTOR does not affect normal presynaptic D2R receptor-dependent motor effects in animals.

**Figure 2.**
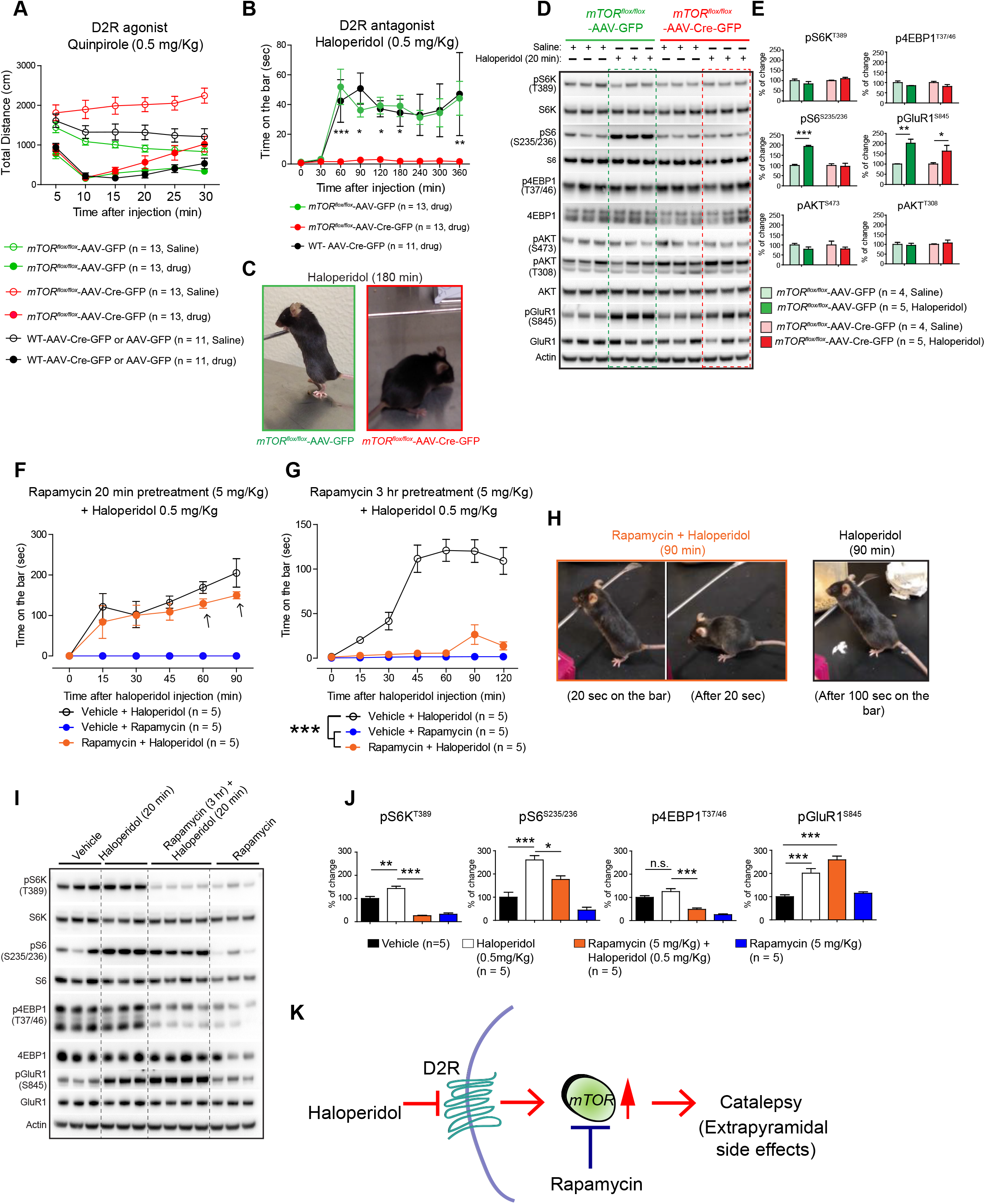
mTOR depletion abolishes D2R antagonist haloperidol-induced catalepsy. (**A**) D2R agonist quinpirole (0.5 mg/Kg, i.p.)-induced open field activity, in AAV-Cre-GFP or AAV-GFP injected *mTOR^foxlflox^* mice and WT control mice. Data are mean ± SEM, n = 11-13 per group, repeated measures two-way ANOVA followed by Bonferroni post-hoc test. (**B**) Catalepsy (as measured by time on the bar)-induced by D2R antagonist haloperidol (0.5 mg/Kg, i.p.) in AAV-Cre-GFP or AAV-GFP injected *mTOR^floxlflox^* and WT control mice. Data are mean ± SEM, n = 1113 per group, *P<0.05, **P < 0.01, ***P < 0.001, repeated measures two-way ANOVA followed by Bonferroni post-hoc test. (**C**) Representative image of AAV-Cre-GFP or AAV-GFP injected *mTOR^flox/flox^* mice treated with haloperidol. (**D**) Western blot analysis of indicated proteins from striatum of indicated mice after 20 minutes of haloperidol (0.5 mg/Kg, i.p.) or saline injection. (**E**) Bar graph indicates quantification of the indicated proteins from C. Data are mean ± SEM, n = 4-5 per group, *P<0.05, **P<0.01, ***P< 0.001, two-way ANOVA, Bonferroni post-hoc test. (**F, G**) Quantification of catalepsy induced by D2R antagonist haloperidol (0.5 mg/Kg, i.p.) in vehicle or pretreated with rapamycin (5 mg/Kg, i.p.) for 20 minutes (**F**) or 3 hours (**G**) in C57BL/6 WT mice. Data are mean ± SEM, n = 5 per group, ***P< 0.001. Repeated measures two-way ANOVA, Bonferroni post-hoc test. (**H**) Representative image of haloperidol induced catalepsy in WT mice pretreated with rapamycin or vehicle. (**I, J**) Western blot analysis (**I**) and quantification (**J**) of indicated targets from the striatal tissue after 20 minutes of haloperidol and rapamycin pretreatment (3 hours). Data are mean ± SEM, n = 5 per group, *P<0.05, **P<0.01, ***P< 0.001. two-way ANOVA, Bonferroni post-hoc test. (**K**) Model shows mTOR mediates D2R inhibitory signals to induce catalepsy linked to extrapyramidal side effects in humans.

Administration of a typical antipsychotic drug (haloperidol (0.5 mg/kg, i.p.), which inhibits post-synaptic D2R (Centonze et al., 2004; Radl et al., 2018; Sebel et al., 2017), robustly induced catalepsy in the control groups (Fig. 2B, C). Conversely, haloperidol administration completely failed to induce any cataleptic effect in *mTOR* mutant mice (Fig. 2B, C). Thus, a striking and complete loss of haloperidol-induced extrapyramidal symptoms was observed in the *mTOR* mutant mice (Fig. 2B, C). These results indicate that *mTOR* depletion interrupts the haloperidol-induced cataleptic effect, suggesting that mTOR signaling selectively controls post-synaptic D2R signaling in the striatal MSNs.

### mTOR mediates haloperidol-induced pS6 phosphorylation in the striatum

Because haloperidol-induced catalepsy is completely abolished in the *mTOR* mutant mice, we hypothesized that haloperidol might promote mTOR signaling in the striatum. We administered haloperidol to *mTOR* mutant mice and the control group and isolated striatal tissue after 20 min. We found a clear upregulation of pS6 (S235/236) by haloperidol only in the AAV-GFP control group but not in the *mTOR* mutant mice (Fig. 2D, E). Surprisingly, haloperidol did not induce the phosphorylation of S6K or the 4EBP1, which are the direct mTORC1 targets. This data is consistent with a previous report(Valjent et al., 2011). Although the reasons for this are unclear, it was proposed that the basal S6K activity may be sufficient to induce pS6 because deletion of S6K abolishes haloperidol-induced pS6 in the striatum(Bonito-Oliva et al., 2013). Phosphoinositide-3 kinase target pAkt (T308) signaling or mTORC2 target pAkt (S473) was also unaltered in the striatum of the treatment and control groups (Fig. 2D, E).

It is well known that by blocking D2R, haloperidol unmasks the ability of adenosine A2AR to enhance striatal cAMP/PKA signaling and ultimately increases the phosphorylation levels of the glutamate receptor subunit GluR1 [pGluR1 (S845)] (Ghiglieri et al., 2015; Valjent et al., 2011). Interestingly, we found that haloperidol robustly induced pGluR1 (S845) in all genotypes and that the extent of activation was similar between *mTOR* mutant and control animals (Fig. 2D, E). Thus, mTOR does not mediate haloperidol-induced pGluR1 signaling in the striatum.

As A2AR and D2R knockout mice showed diminished haloperidol-induced catalepsy (Boulay et al., 2000; El Yacoubi et al., 2001) and because haloperidol acts by blocking of D2R, we wanted to confirm that striatal expression of these receptors was comparable in the *mTOR* mutant mice and control mice. We found similar A2AR levels (but significantly enhanced D2R levels) in the striatum of the *mTOR* mutant mice compared to the control (Supplementary Figure 1). Thus, diminished haloperidol-induced catalepsy is not due to diminished A2AR or D2R levels in the *mTOR* mutant mice.

### Pharmacological inhibition of mTOR abolishes haloperidol-induced catalepsy

As striatal genetic depletion of mTOR completely abolished the haloperidol-induced catalepsy, we next asked whether pharmacological inhibition of mTOR would produce a similar phenotype. To investigate this, we treated 4-month-old C57BL/6 WT mice with mTORC1 inhibitor rapamycin (5.0 mg/kg., i.p.) for 20 minutes, followed by injection with haloperidol (0.5 mg/kg, i.p.). Haloperidol promoted a time-dependent cataleptic behavioral response in the vehicle-injected C57BL/6 mice, as well as in the rapamycin pretreated C57BL/6 WT mice (Fig. 2F). As expected, rapamycin treatment alone did not elicit a catalepsy response (Fig. 2F). This result indicates that 20 min of pretreatment with rapamycin does not affect the haloperidol-induced cataleptic response.

Interestingly, although the onset of the cataleptic behavioral response was similar between the groups, there was a trend towards decreased cataleptic behavior in the rapamycin pretreated animals after 60 and 90 min post haloperidol administration (Fig. 2F, arrow). This observation prompted us to hypothesize that rapamycin may interfere with a cataleptic response after 60 min or longer duration following administration. To investigate this hypothesis, we pretreated C57BL/6 WT mice with rapamycin (5.0 mg/kg., i.p.) for 3 hours before administering haloperidol (0.5 mg/kg, i.p.). Strikingly, we found a dramatic attenuation of haloperidol-induced catalepsy in animals that were pretreated with rapamycin for 3 hours (as compared to vehicle-treated groups) (Fig. 2G, H). This result indicates that a more prolonged exposure to rapamycin [which may be necessary for target (mTOR) engagement] is a prerequisite to block the haloperidol-induced cataleptic response in mice.

### Pharmacological inhibition of mTOR diminishes haloperidol-induced pS6 but not pGluR1

Next, we investigated how rapamycin pretreatment impacted on haloperidol-induced striatal signaling in C57BL/6 WT mice. Compared to the vehicle, we found that haloperidol (for 20 min) robustly induced striatal pS6 (S235/236) and pGluR1 (S845) signaling in C57BL/6 WT mice (Fig. 2I, J). Haloperidol did not induce p4EBP1 (T37/46) in C57BL/6 WT mice, consistent with genetic model (Fig. 2D, E). However, a slight but significant increase of pS6K (T389) (Fig. 2I, J) was observed in haloperidol-injected C57BL/6 WT mice. Rapamycin pretreatment suppressed the haloperidol-induced pS6K and pS6 as well as diminished the basal pS6K, pS6, and p4EBP1. Rapamycin did not interfere with pGluR1 signaling, in the striatum (Fig. 2I, J), consistent with the observation in genetic model (Fig. 2D, E). These results indicate that pharmacological blocking of mTORC1 by rapamycin prevents the haloperidol-mediated mTORC1 signaling and associated catalepsy in the striatum.

## DISCUSSION

The data presented here indicate that mTOR signaling in the striatum mediates post-synaptic D2R-mediated functions, as both genetic depletion of mTOR or pharmacological inhibition of mTORC1 signaling by rapamycin prevented a haloperidol-induced catalepsy response (Fig. 2K). Importantly, mTOR regulates specific signaling and behavioral functions in the striatum. The D1R-mediated motor behaviors and the presynaptic D2R signaling are unaffected by the loss of striatal mTOR. Our data represent, to the best of our knowledge, the first report to use rapamycin to assess the role of mTOR signaling in haloperidol-induced catalepsy.

Previous studies showed that haloperidol induces pS6 signaling by enhancement of adenosine A2A/Golf signaling; however, the functional relevance of this pathway and its role in cataleptic behaviors were unknown (Bowling et al., 2014; Valjent et al., 2011). PKA signaling that induces pGluR1 is particularly linked to the generation of haloperidol-induced catalepsy. (Adams et al., 1997; Roche et al., 1996). Studies in non-neuronal cells showed that PKA acts upstream of mTOR and can activate or inhibit it (de Joussineau et al., 2014; Jewell et al., 2019; Kim et al., 2010). PKA can directly phosphorylate mTOR and promote the phosphorylation of S6K in adipose tissue. (Liu et al., 2016). Indeed, it has been demonstrated that PKA activation induces pS6 in cultured striatal neurons (Valjent et al., 2011). With rapamycin or mTOR depletion, we found that mTOR signaling in the striatum did not interfere with haloperidol-induced pGluR1 signaling; however, it altogether abolished haloperidol-induced catalepsy. Thus, our data indicate that mTOR signaling in the striatum promotes haloperidol-induced catalepsy by acting downstream or independently of PKA-pGluR1 signaling.

The results presented here clearly suggest that an acute pretreatment of rapamycin completely reverses haloperidol-induced catalepsy, further emphasizing the critical role of mTORC1 in altering D2R signaling to promote extrapyramidal symptoms. Note that short-term (20 min) pretreatment with rapamycin had a negligible effect. However, long-term rapamycin pretreatment (3 hours) abolished haloperidol-induced catalepsy. One possibility for such delayed action is due to the relatively poor brain penetrability of rapamycin and thus a delayed target engagement (Brandt et al., 2018). Interestingly, a previous study indicated Fyn kinase had a role in the regulation of haloperidol-induced catalepsy (Hattori et al., 2006). Fyn kinase also promotes mTORC1 signaling and it is therefore tempting to speculate that Fyn-mTORC1 signaling may have a role in haloperidol-mediated catalepsy (Hattori et al., 2006; Wang et al., 2015).

What are the molecular mechanisms underlying haloperidol–mTORC1–cataleptic behavior? A previous study indicated that haloperidol induced the mTOR-dependent translation and neuronal morphology in cultured MSNs (Bowling and Santini, 2016). In vivo, haloperidol can increase or decrease MSN morphology (spine density); in particular, it can decrease the spine density in D2 MSN (Sebel et al., 2017). Therefore, it is conceivable that mTOR is a critical regulator of haloperidol-induced molecular changes in the striatum. In addition to protein synthesis, mTOR signaling also regulates autophagy, purine, and lipid biosynthesis (Ben-Sahra and Manning, 2017). Based on our study, it is possible that mTOR signaling may translate the haloperidol-induced signaling into catalepsy by more than one pathway. Further research on the importance of these mechanistic insights, by dissecting the cell-type-specific role of mTOR, identification of haloperidol-induced mTOR interactors, and high-throughput comparative proteomic analysis in mTOR mutant and WT mice, could help unravel D2R-specific mechanisms of mTOR signaling in extrapyramidal symptoms.

Haloperidol is a major antipsychotic medication prescribed to diminish psychosis in schizophrenia patients (Ostinelli et al., 2017). However, its action is limited due to its elicitation of Parkinsonian-like bradykinesia, which affects a majority of patients and is commonly called haloperidol-induced EPS (Finucane et al., 2020; Kurz et al., 1995). To date, there are no effective treatments available for haloperidol-induced EPS. By combining genetic and pharmacological approaches, our mechanistic models provide a clear insight into the causal role of mTOR signaling in promoting haloperidol-induced catalepsy in preclinical murine systems (Fig. 2K). Our study therefore illustrates the translational potential of rapamycin in alleviating striatal D2R-mediated EPS in humans.

## Author contributions

S.Su conceptualized the project. UNRJ designed and carried out all the work in *mTOR^flox/flox^* and related control mice. A.U directed, and N.S co-designed, pharmacological experiments with UNRJ. W.P performed preliminary behavioral analysis. S.Su wrote the paper with input from the co-authors.

## Acknowledgments

We would like to thank Melissa Benilous for her administrative help and the members of the lab for their continuous support and collaborative atmosphere. This research was supported by funding from National Institutes of Health/National Institute of Neurological Disorders and Stroke grant R01-NS087019-01A1, National Institutes of Health/National Institute of Neurological Disorders and Stroke grant R01-NS094577-01A1 and a grant from Cure Huntington Disease Initiative (CHDI) foundation.

## Disclosure Statement

The authors have no conflicts of interest to disclose.

---

**Supplementary Figure 1.**
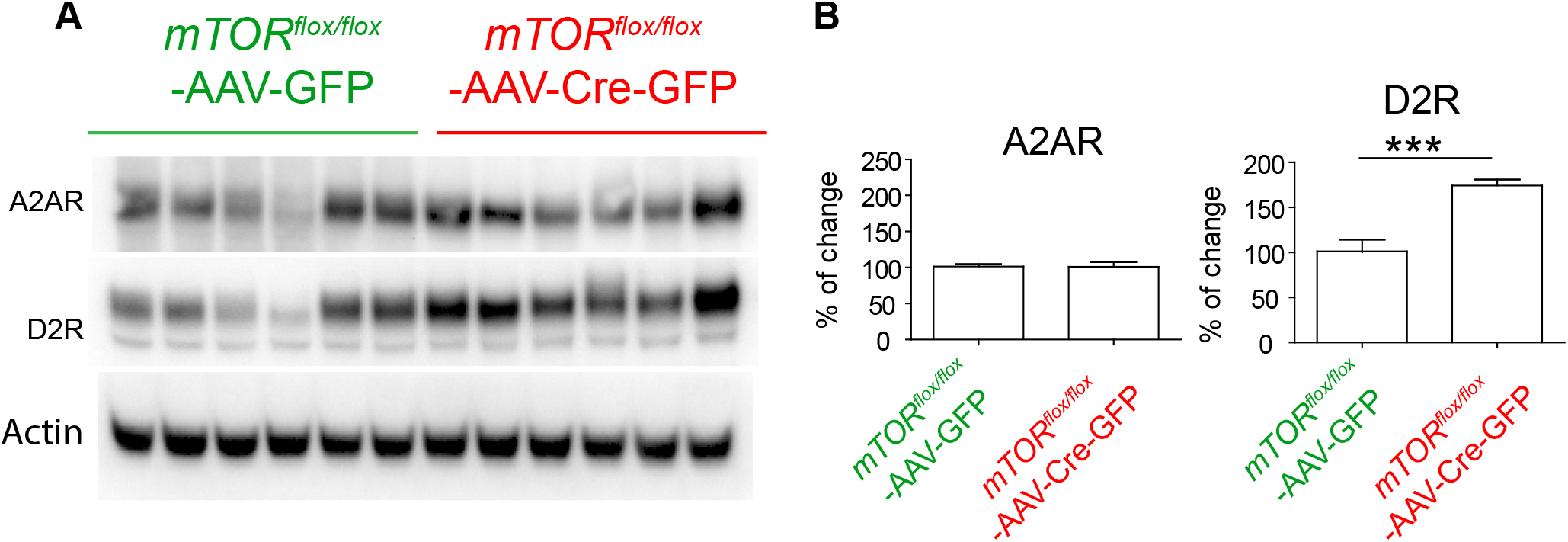
A2AR and D2R expression in *mTOR* mutant mice. Western blot analysis of indicated proteins from striatum of AAV-GFP and AAV-Cre-GFP injected *mTOR^flox/flox^* mice. (**B**) Bar graph indicates quantification of the indicated proteins from A. Data are mean ± SEM, n = 4-5 per group, ***P< 0.001. two-way ANOVA, Bonferroni post-hoc test.

## MATERIALS AND METHODS

### Chemicals and Antibodies

The majority of the chemicals used were purchased from Sigma, unless mentioned otherwise. Antibodies against - mTOR (#2983) pS6K T389 (#9234), S6K (#9202), pS6 S235/236 (#4858), S6 (#2217), p4EBP1 T37/46 (#2855), 4EBP1 (#9644), pAkt S473 (#4060), pAkt T308 (#13038), and Akt (#4691) were from Cell Signaling Technology. Antibodies for actin (sc-47778), GFP (sc-33673), A2AR (sc-32261), and D2R (sc-5303) were from Santa-Cruz Biotechnology. Ctip2 (ab18465) antibody was from Abcam. (+/-)-Quinpirole dihydrochloride, (Q111), Haloperidol (H1512), and R(+)-SCH-23390 hydrochloride (D054) were purchased from MilliporeSigma. SKF 81297 hydrobromide was from R&D systems. Rapamycin was purchased from LC laboratories (R-5000). Haloperidol was initially dissolved in glacial acetic acid, then its pH was adjusted close to 7 with NaOH, and final dilution was made in saline solution (0.9 %). Rapamycin was dissolved in 5% dimethyl sulfoxide (DMSO), 15% PEG-400 (polyethylene glycol, molecular weight 400), and 5% Tween-20, and finally dissolved in saline solution for injection. SCH23390, SKF81297, and Quinpirole were dissolved in saline solution. All the drugs were administrated by intraperitoneally (i.p.) injection.

### Animals

*mTOR^flox/flox^* mice that harbor *loxP* sites flanking exons 1-5 of the mTOR locus (The Jackson Laboratory, strain *B6.129S4-mTOR^tm1.2Koz^*/J, Stock No: 011009) and C57BL/6J (wild type, WT) were used for adeno-associated virus micro-injections. Mice were housed in groups of two or three on a 12:12 h light-dark cycle and were provided food and water ad libitum. All protocols were approved by Institutional Animal Care and Use Committee at The Scripps Research Institute, Florida,

### Stereotaxic surgeries

For all surgical procedures, eight-week old mice were anesthetized through the constant delivery of isoflurane while mounted in a stereotaxic frame (David Kopf Instruments). AAV-Cre-GFP (AAV1.hSyn.HI.eGFP-Cre.WPRE.SV40) or AAV-GFP (green fluorescent protein, AAV1.hSyn.eGFP.WPRE.bGH) (Vector Core, University of Pennsylvania) were injected bilaterally into the striatum at the following coordinates: ML = ± 1.6, AP = +1.1; DV = −3.9/−3.5 and ML = ± 2.5; AP = +0.5; and DV = −4.2/−3.6 from bregma. Virus was injected in 0.5 μl volumes (5.9X10^12^ gc/mL) per injection site in each animal (4 μl total). Animals recovered for two weeks before behavioral testing. The efficacy of the viral injections was determined by GFP expression in the striatum.

### Behavioral Analysis

Longitudinal behavioral testing was performed for AAV-Cre-GFP or AAV-GFP-injected *mTOR^flox/flox^* mice, and wild type (WT) mice. All behavioral testing was performed as described in our previous work(Pryor et al., 2014; Swarnkar et al., 2015) during the light phase of the light-dark cycle between 8:00 am and 12:00 pm. For each week/month of behavioral testing, the following measures were assessed with the rotarod test on the first four days and an open-field test on the fifth. Rotarod testing was performed using a linear accelerating rotation paradigm (Med Associates Inc.) for three trials separated by 20 min for four consecutive days each month. The mice were placed on the apparatus at 4 rpm and were subjected to increasing rpm, accelerating to 40 rpm over the course of a maximum of five minutes. The overall latency to fall for each mouse was calculated as the average of the three trials across four days for each month. The latency of falling from the rod was scored as an index of motor coordination, while improvement in performance across training days, as measured by increasing latency to fall from the rotarod, indicates motor learning. Open-field activity was assessed in a single 30-minute session using EthoVision XT software (Noldus Information Technology). Each mouse was placed individually in the center of each square enclosure, and movement was quantified automatically. Single cohort of mice with mixed sex ratio were used for the behavior testing of AAV injected *mTOR^flox/flox^* mice: *mTOR^flox/flox^*-AAV-GFP (n = 13, male 3 and female 10) and *mTOR^flox/flox^*-AAV-Cre-GFP (n = 13, male 7 and female 6). Single cohort of mice were used for behavior testing of AAV injected WT mice: WT-AAV-GFP (n = 5, female 2, male 3) and WT-AAV-Cre-GFP (n = 6, female 3, male 3). As there were no differences in the behaviors of WT-GFP and WT-Cre-GFP injected mice, they were combined for the group analysis.

#### Quinpirole and SKF81297 evaluation

Pharmacological effect of D2R agonist Quinpirole and D1R agonist SKF81297 was made using the same Open-field system and the EthoVision XT software (Noldus Information Technology).

Each mouse was placed individually in the center of a plastic box (11x 14 inches) with fresh bedding. For SKF81297 evaluation, mice were placed in the boxes for 30 minutes, as basal activity and habituation, then the drug was injected (2.5 mg/Kg, i.p.) and the total activity was recorded for 90 minutes. Results were plotted in a bar-graph showing the % change in the total activity after habituation at 0, 30 and 60 minutes. For Quinpirole experiment, mice were placed in the center of the plastic box with fresh bedding after the drug injection (0.5 mg/Kg, i.p.), then the total distance traveled (cm) was measured each 5 min during the next 30 min. Before each protocol, animals were kept in a waiting room for at least 30 minutes. Each control group was treated with the vehicle according to the drug.

#### Haloperidal and SCH23390 evaluation

Behavioral evaluation for D2R antagonist Haloperidol (0.5 mg/Kg, i.p.) and D1R antagonist SCH23390 (0.1 mg/Kg, i.p.) was made by measuring the catalepsy-induced effect using the bar test. Catalepsy was determinate placing each mouse with its forelegs on the bar in a kangaroo posture (Figure1L and Figure 2C, H), latency to change the corporal posture was recorded for three trials, and average of them was used for group analysis. After drug injection mice were evaluated on the bar at 15, 30, 60, 90 for SCH23390, and at 0, 30, 60, 90, 120, 180, 240, 300, and 360 minutes for Haloperidol in the *mTOR^flox/flox^* and WT mice injected with AAV-GFP or AAV-Cre-GFP viruses. For Rapamycin and Haloperidol experiments in C57BL/6 WT mice, animals were injected with Rapamycin (5 mg/Kg) as pretreatment at 20 minutes or 3 hours before Haloperidol (0.5 mg/Kg). After Haloperidol injection mice were tested on the bar at 15, 30, 45, 60, 90, and 120 minutes. Before each protocol, each mouse was kept in single cage in the procedure room for at least 30 minutes. Each control group was treated with the vehicle according to the drug.

### Western blot analysis

Twenty minutes after haloperidol injection, mice were euthanized by decapitation and brains were rapidly dissected and the striatum was quickly removed and snap-frozen in liquid nitrogen. Tissue was homogenized in RIPA buffer [50 mM Tris-HCl (pH 7.4), 150 mM NaCl, 1.0% Triton X-100, 0.5% sodium deoxycholate, 0.1% SDS,) with a protease inhibitor cocktail (Roche, Sigma) and phosphatase inhibitors (PhosSTOP, Roche, Sigma). Protein concentration was measured using BCA protein assay reagent (Pierce). Protein lysates were loaded and separated by 4-12% Bis-Tris Gel (Invitrogen), transferred to PVDF membranes, and probed with the indicated antibodies. Secondary antibodies were HRP-conjugated (Jackson Immuno Research, Inc).

Chemiluminescence was detected using WesternBright Quantum (Advansta) ECL reagent using a chemiluminescence imager (Alpha Innotech). Western blotting experiment was carried out as described previously(Pryor et al., 2014; Shahani et al., 2017; Shahani et al., 2014; Shahani et al., 2016; Swarnkar et al., 2015). Relative levels of all the proteins were normalized to actin and quantified using Image J. Relative levels of phosphorylated proteins were normalized to respective normalized total proteins and quantified.

### Immunohistochemistry and analysis

Immunostaining was performed as previously described (Chen et al., 2015; Shahani et al., 2017; Swarnkar et al., 2015). Briefly, mouse brains were fixed in 4% paraformaldehyde for overnight, cryoprotected in a sucrose/PBS gradient at 4 °C (10, 20 and 30%), and embedded in Tissue-Tek OCT compound (Sakura). Coronal sections (20μm) were collected on Superfrost/Plus slides and immunostained after heat-induced antigen retrieval [10 min in boiling citrate buffer (pH 6.0), MilliporeSigma, C9999]. Primary antibodies used in this study were anti-Ctip2 (1:500, Abcam, ab18465), anti-mTOR (1:250, Cell Signaling, #2983), and anti-GFP (Santa Cruz, SC33673). Alexa Fluor 488, 594, and 647 conjugated secondary antibodies (Thermo Fisher Scientific) were used in this study. Immunofluorescent brain sections were counterstained with DAPI and mounted using Fluoromount-G mounting medium (Thermo Fisher Scientific). Images were obtained with the Zeiss LSM 880 microscope and processed using the ZEN software (Zeiss).

For cell quantification, five regions of interest (ROIs) of 100 μm^2^ were defined in immunostained sections (four to five sections for each mouse, n= 4 mice per group) of the medial striatum from *mTOR^flox/flox^* injected with AAV-GFP or AAV-Cre viruses. Total number of cells were calculated by counting the DAPI stained nuclei. AAV-Infected neurons were identifying by expression of the GFP. GFP expression was observed in the soma of the AAV-GFP infected neurons while AAV-Cre infected neurons expressed GFP in the nucleus. Percentage of the Ctip2, mTOR and GFP triple-positive neurons were determined considering DAPI stained cells as 100%. Ventricular area was determinate in hematoxylin/eosin-stained sections, from the same animals, four rostral (+1.1 from Bregma) and caudal (+0.5 from Bregma) sections from each animal (n=4 mice per group) were taken using the Leica DM5500B microscope. The ventricular area was calculated by analyzing the images using the ImageJ software.

### Statistical Analysis

Data are presented as mean ± SEM as indicated. Statistical analysis was performed with a Student’s t-test or two-way ANOVA followed by Bonferroni post-hoc test or repeated measure two-way ANOVA followed by Bonferroni post-hoc test as indicated in the figure legends. Repeated measures two-way analysis of variance (ANOVA) where time was the repeated measure and treatment/genotype group was the fixed effect. Post hoc Bonferroni multiple comparison tests were used to identify statistically significant differences between treatment/genotype groups at each time point. Significance was set at P < 0.05. All statistical tests were performed using Prism 7.0 (GraphPad software).

## Notes

### Competing Interest Statement

The authors have declared no competing interest.

